# High-fat and high-sucrose diet-induced hypothalamic inflammation shows sex specific features in mice

**DOI:** 10.1101/2024.07.29.605582

**Authors:** Gabriela C. De Paula, Rui F. Simões, Alba M. Garcia-Serrano, João M. N. Duarte

**Affiliations:** Diabetes and Brain Function Unit, Department of Experimental Medical Science, Faculty of Medicine, Lund University, Lund, Sweden; Wallenberg Centre for Molecular Medicine, Faculty of Medicine, Lund University, Lund, Sweden; Institute for Research in Biomedicine, Bellinzona, Switzerland

**Keywords:** neuroinflammation, high fat, sucrose, reverse diet, cytokines, gliosis

## Abstract

Hypothalamic inflammation underlies diet-induced obesity and diabetes in rodent models. While diet normalization largely allows for recovery from metabolic impairment, it remains unknown whether long-term hypothalamic inflammation induced by obesogenic diets is a reversible process. In this study, we aimed at determining sex specificity of hypothalamic neuroinflammation and gliosis in mice fed a fat- and sugar-rich diet, and their reversibility upon diet normalization. Mice were fed a 60%-fat diet complemented by a 20% sucrose drink (HFHSD) for 3 days or 24 weeks, followed by a third group that had their diet normalized for the last 8 weeks of the study (reverse diet group, RevD). We determined the expression of pro- and anti-inflammatory cytokines, and of the inflammatory cell markers IBA1, CD68, GFAP and EMR1 in the hypothalamus, and analyzed morphology of microglia (IBA-1^+^ cells) and astrocytes (GFAP^+^ cells) in the arcuate nucleus. After 3 days of HFHSD feeding, male mice showed over-expression of IL-13, IL-18, IFN-γ, CD68 and EMR1 and reduced expression of IL-10, while females showed increased IL-6 and IBA1 and reduced IL-13, compared to controls. After 24 weeks of HFHSD exposure, male mice showed a general depression in the expression of cytokines, with prominent reduction of TNF-α, IL-6 and IL-13, but increased TGF-β, while female mice showed over-expression of IFN-γ and IL-18. Furthermore, both female and male mice showed some degree of gliosis after HFHSD feeding for 24 weeks. In mice of both sexes, diet normalization after prolonged HFHSD feeding resulted in partial neuroinflammation recovery in the hypothalamus, but gliosis was only recovered in females. In sum, HFHSD-fed mice display sex-specific inflammatory processes in the hypothalamus that are not fully reversible after diet normalization.

## INTRODUCTION

Obesity is nowadays considered a 21^st^ century pandemic and has emerged one of the major health risk factors with economic burden [1, 2]. This condition can be defined as excessive adipose tissue accumulation due to an imbalance between energy intake and expenditure [3, 4]. While in genetic obesity a mutation in a gene deregulates energy homeostasis, environmental obesity can be caused by the ingestion of obesogenic diets that are highly caloric, and rich in lipids and sugar [5-7]. The prevalence of obesity is known to be sex-specific, being women more affected than men [1, 2, 8]. This can be due, in part, to the inherent biological difference between the two sexes, in which females have higher body fat proportion compared to males [2]. Sex hormones and menopause also play key roles on obesity development. It has been described that low levels of testosterone can be associated with obesity since this hormone was shown to promote fat consumption [2, 9, 10].

Obesity has been known to induce systemic inflammation that predisposes individuals to the development of comorbidities, including cardiovascular diseases, metabolic syndrome and type 2 diabetes, conditions which in turn impact the brain [11-13]. We have previously shown that long-term exposure to an obesogenic diet rich in saturated fat and sucrose induced reversible alterations in cortex and hippocampus function (behavior) and metabolism in mice [14]. Increased activation of microglia, the brain resident immune cells, was found in the same brain regions, without over-expression of pro-inflammatory cytokines [14], which suggests a sustained low-grade neuroinflammation. In this setting, any cortical and hippocampal alterations were normalized after diet reversal to a low-fat and low-sugar diet [14].

There is increasing evidence that the innate immune activation in the hypothalamus is key element in the pathogenesis of diet-induced obesity. The hypothalamus is the central regulator of body weight and energy homeostasis, integrating and controlling nutrient-sensing signals [15-17]. Hypothalamic inflammation, induced by obesogenic diets, leads to the alteration of normal hypothalamic function that impact feeding behavior and the balance between energy intake and expenditure [18, 19]. Additionally, diet-induced hypothalamic inflammation is typically characterized by the activation of reactive astrocytic and microglial cells, with pro-inflammatory cytokines burden and the subsequent inflammatory cascade [18, 20]. Previous studies using murine models fed high-fat diets (HFD) or high-fat and high-sucrose diets (HFHSD) for different timepoints evidenced this link between inflammation and brain disfunction. A study using blood oxygen level-dependent (BOLD) functional magnetic resonance imaging (fMRI) on mice fed a HFHSD for only 7 days clearly showed hypothalamic dysfunction, namely impaired response to glucose administration [21]. Moreover, HFD-fed mice exhibited an increase in typical astrogliosis and microgliosis markers after 2, 4 and 6 months, including increased density of GFAP and IBA1, and number of GFAP^+^ and IBA1^+^ cells and/or their area [22, 23].

The above-mentioned studies focused on male rodents [18, 20-23]. Male mice fed obesogenic diets are the model of election for metabolic syndrome development. That is because sex-differences on the response to dietary fat are well documented, especially the limited development of insulin sensitivity and hyperinsulinemia in female mice [14, 24]. Thus, little is known on how obesity and metabolic syndrome development during HFD or HFHSD exposure impacts the brain of female rodents, in particular in the hypothalamus. Hypothalamic inflammation differences have been recently reported by Church *et al*. [25] and Daly *et al*. [26]. An earlier study has also proposed differential inflammatory profiles in the hippocampus of male and female mice born from HFD-fed dams [27]. These studies further suggest sex differences in the interaction between gut and brain inflammation. In our previous work, the cortex and hippocampus of mice of either sex showed recovery of HFHSD-induced alterations upon diet normalization [14], but it is hitherto unknown whether reversibility of neuroinflammation occurs in the hypothalamus. Using a cohort of mice from this previously reported study [14], we now determined if hypothalamic inflammation induced by HFHSD is a sex-specific reversible process.

## MATERIALS AND METHODS

### Animals

Experiments were performed according to EU Directive 2010/63/EU, approved by the Malmö/Lund Committee for Animal Experiment Ethics (#994/2018), and are reported following the ARRIVE guidelines (Animal Research: Reporting in Vivo Experiments, NC3Rs initiative, UK). 8-weeks old male and female C57BL/6J mice (RRID:IMSR_JAX:000664) were purchased from Taconic Biosciences (Köln, Germany), and housed in groups of 3-5 animals on a 12h light-dark cycle with lights on at 07:00, room temperature of 21-23 °C and humidity at 55-60%. Mice were habituated to the facility for 1 week upon arrival and subsequently randomly assigned to 5 experimental groups, receiving either a 10%-fat control diet (CD) for 3 days or 24 weeks, a composition-matched high-fat diet (60%-fat) plus access to a 20%(w/v) sucrose in drinking water (HFHSD), for 3 days or 24 weeks, or a reversed diet, that consisted of HFHSD feeding for 16 weeks followed by CD feeding for 8 weeks (figure 1A-B). Mice receiving HFHSD also had access to sugar-free water. Food and water were provided *ad libitum*.

**FIGURE 1.**
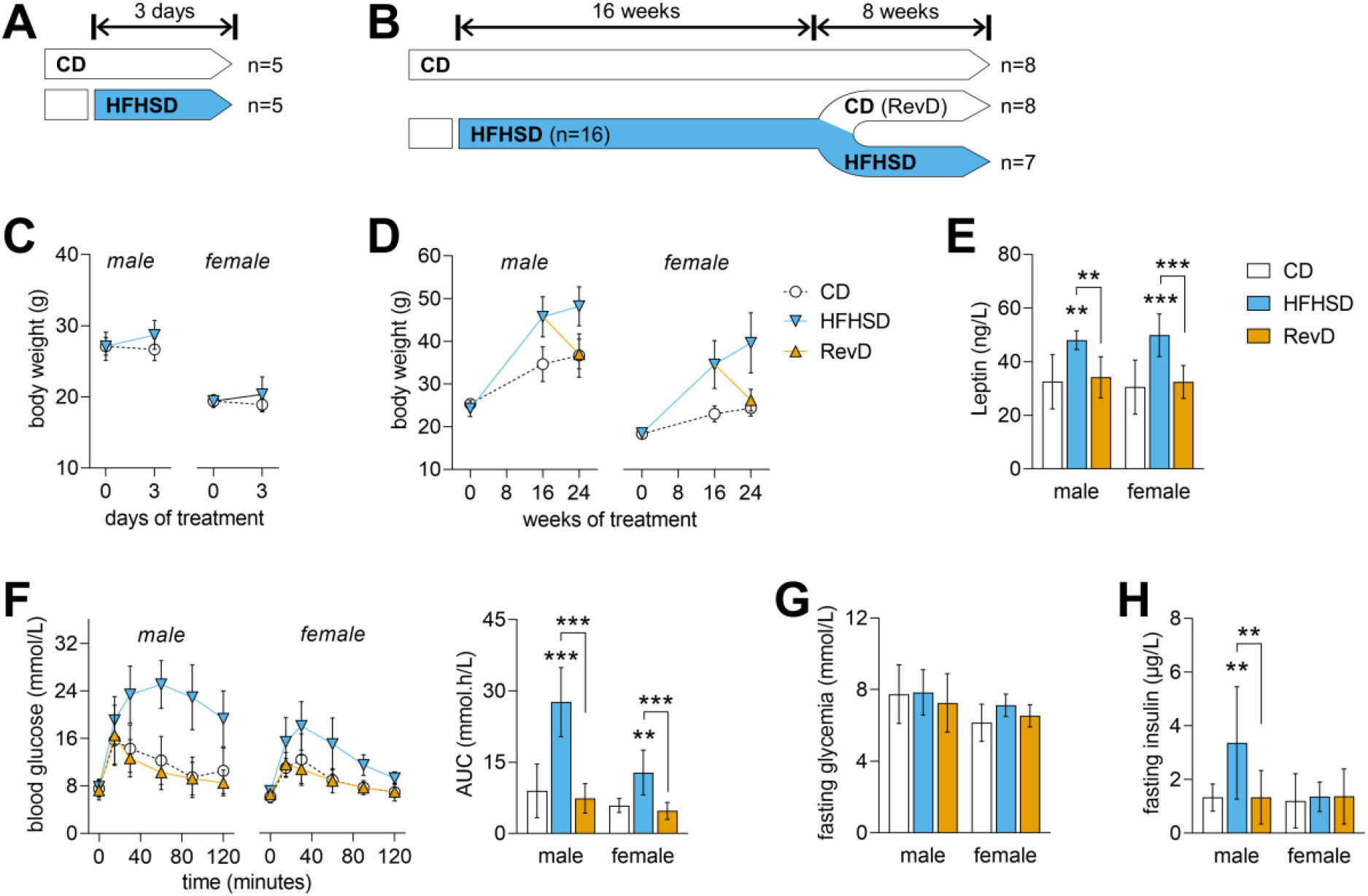
Experimental study design and number of mice (n) per sex in the experimental group (A-B), body weight (C-D), and metabolic assessments after 24 weeks of dietary intervention (E-H). Body weight of male and female mice shows obesity development upon HFHSD feeding, and recovery after diet normalization (D). Plasma leptin levels after 24 weeks of dietary intervention is increased by HFHSD and normalised in RevD (E). Glucose clearance in GTT was reduced by HFHSD feeding, and the area under the curve (AUC) of the GTT shows full recovery after diet normalization (F). Fasting glycemia (G) and plasma insulin levels (H) are indicative of HFHSD-induced insulin resistance in males but not females. Data shown as mean±SD of n=5-as depicted in (A-B); **P<0.01, ***P<0.001 for comparing HFHSD *versus* CD or as indicated.

Diets were acquired from Research Diets (New Brunswick, NJ-USA): a high-fat lard-based diet with 60% kcal of fat (D12492) and a control diet containing 10% kcal of fat (D12450J), with total energy of 5.21 and 3.82 kcal/g, respectively [14].

After 24 weeks on HFHSD, we carried out glucose tolerance tests (GTT) and plasma hormone analyses as detailed previously [14]. Brain samples were collected at 3 days or 24 weeks of diet intervention after decapitation under brief isoflurane anesthesia.

### Real-time polymerase chain reaction (RT-PCR)

RNA was isolated from mice hypothalamus with Trizol (#15596026, Invitrogen, USA), and reverse transcribed using the script cDNA SuperMix (#95048, Quantabio, England), and then the resulting cDNA was used for quantitative RT-PCR as detailed by [28] using PerfeCTa SYBRgreen SuperMix (#95054, Quantabio, England) and primers for allograft inflammatory factor 1 (IBA1), cluster of differentiation 68 (CD68), adhesion G protein-coupled receptor E1 (EMR1), glial fibrillary acidic protein (GFAP), interleukin (IL)-1β, IL-6, IL-10, IL-13, IL-18, interferon γ (IFN-γ), transforming growth factor β (TGF-β), tumor necrosis factor α (TNF-α), and β-actin (Table 1). Gene expression was normalized to β-actin expression with the comparative cycle threshold method (ΔΔCT).

**TABLE 1.**
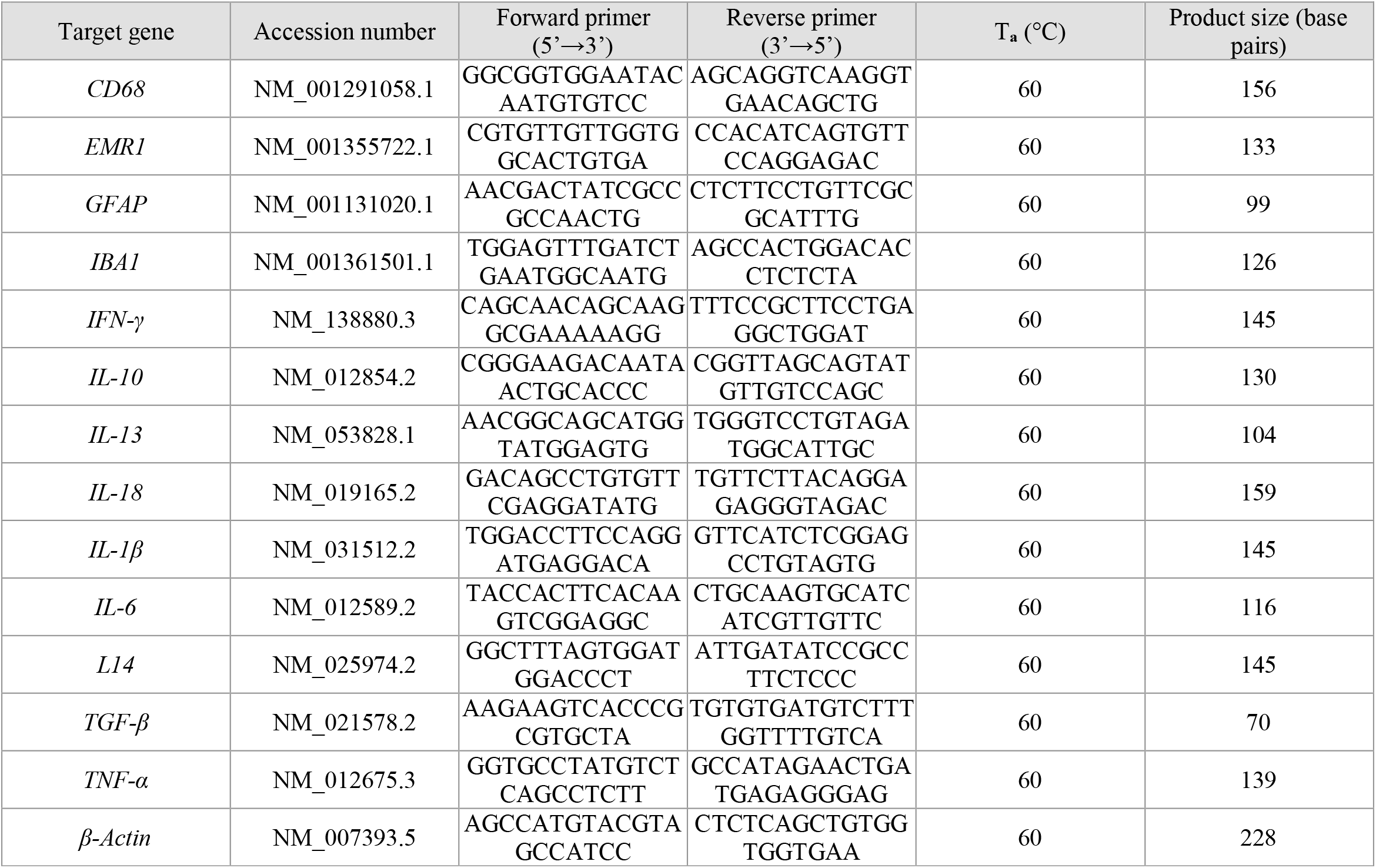
Primers used for real time-PCR, respective gene, accession number, annealing temperature (T_a_), and measured primer size.

### Immunofluorescence confocal microscopy

Mice were sacrificed under isoflurane anesthesia by cardiac perfusion with cold PBS and then cold phosphate-buffered formaldehyde (Histolab, Askim, Sweden), and brains were cryosectioned into 20 µm slices [29]. Immunolabeling was carried out as detailed previously [21] with the primary antibodies: rabbit anti-allograft inflammatory factor 1 (IBA1, dilution 1:200; #019-19741, Fujifilm Wako, Japan; RRID:AB_839504), and anti-glial fibrillary acidic protein (GFAP) pre-tagged with AF488 (dilution 1:500; #53-9892-82, ThermoFisher Scientific, Göteborg, Sweden; RRID:AB_10598515). Secondary antibody (dilution 1:500) was from ThermoFisher: AF568-conjugated goat anti-Rabbit IgG (#A-21069; RRID: AB_141416). After mounting, slices were examined under a Nikon A1RHD confocal microscope (Nikon Instruments, Tokyo, Japan). Images were acquired with NIS-element v5.20.01 (Laboratory Imaging, Nikon), and analyzed in ImageJ (NIH, Bethesda, MD, USA; RRID:SCR_003070) as previously [14].

### Statistics

Results are presented as mean±SD unless otherwise stated. Partial least-squares (PLS) regression with 5 components was applied on z-scores of gene expression profiles using MATLAB 2019a (MathWorks, Natick, MA-USA; RRID:SCR_001622). The PLS model was fit to each CD-HFHSD paired dataset, and the variable importance in projection (VIP) was calculated for each gene. Data from the RevD group was not analyzed but reconstructed with the obtained models for male and female mice. One-sample t-tests were used for determining specific gene expression changes. Prism 9.4.0 (GraphPad, San Diego, CA-US; RRID:SCR_002798) was used for analysis of metabolic phenotype and immunofluorescence results. After assessing normality with the Kolmogorov-Smirnov and Shapiro-Wilk tests, data were analyzed with the two-way ANOVA followed by independent comparisons with the Fisher’s least significant difference (LSD) test. Statistically significant differences were considered for P<0.05.

## RESULTS

As reported previously, HFHSD feeding induces obesity with glucose intolerance in both genders, and hyperinsulinemia in male but not female mice [14]. In particular, 3 days of HFHSD exposure was sufficient to induce a small weight increase (figure 1C; weight gain ANOVA: gender P=0.396, diet P<0.001, interaction P=0.533), which is pronounced after 24 weeks, and recovers upon diet normalization (figure 1D; weight gain ANOVA: gender P<0.001, diet P<0.001, interaction P<0.001). In line with obesity and increased fat deposition [14], HFHSD but not RevD mice showed increased fed leptin in plasma (figure 1E; ANOVA: gender P=0.820, diet P<0.001, interaction P=0.800). Male mice fed the HFHSD showed more severe glucose intolerance than females in a GTT (figure 1F; ANOVA: gender P<0.001, diet P<0.001, interaction P<0.001). HFHSD had negligible effects on fasting glycaemia, although females showed lower blood glucose than males (figure 1G; ANOVA: gender P=0.006, diet P=0.330, interaction P=0.510), while fasting insulinaemia was impacted by HFHSD in males but not females (figure 1H; gender P=0.029, diet P=0.042, interaction P=0.018). Finally, diet normalization after 16 weeks of HFHSD exposure resulted in recovery of body weight, glucose tolerance, and hyperinsulinemia (in males) to control values (figure 1).

### Effect of HFHSD-feeding for 3 days on hypothalamic inflammation markers

Thaler *et al*. reported hypothalamic inflammation within a few days after HFD exposure in mice [18]. Thus, we first analyzed mRNA expression levels of pro-inflammatory cytokines IL-1β, IL-6, IL-18, TNF-α and IFN-γ, anti-inflammatory cytokines IL-10, IL-13 and TGF-β, as well as astrogliosis (GFAP) and microgliosis (IBA1, CD68 and EMR1) markers in the hypothalamus of male and female mice after 3 days HFHSD feeding. A PLS regression with 5 components provided a separation of mice in HFHSD or CD based on their gene expression (figure 2) and explained 98% and 95% of the variance in males and females, respectively.

**FIGURE 2.**
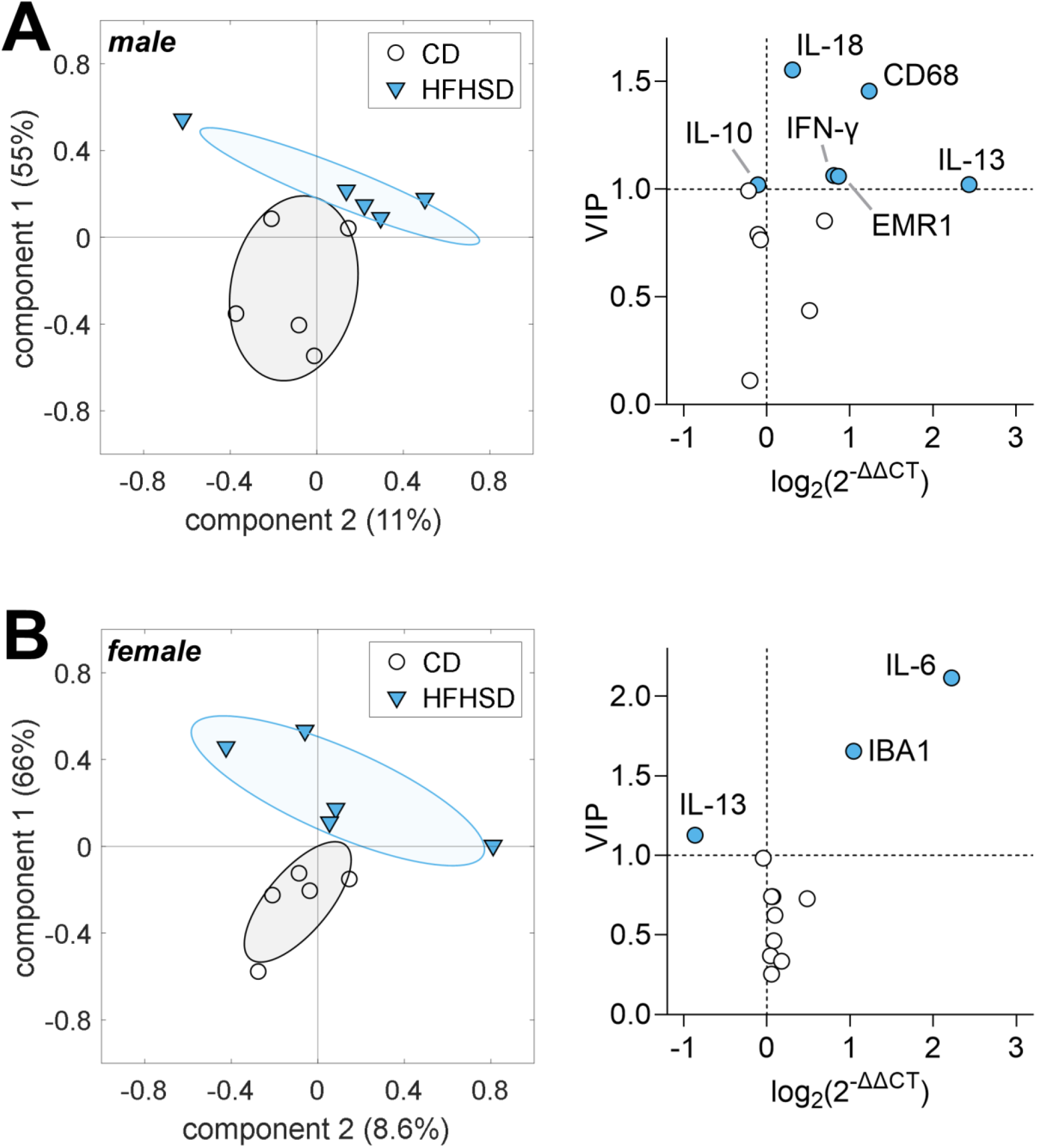
HFHSD feeding for 3 days induced gender-specific hypothalamic inflammation, that is, differential gene expression changes were observed in male (A) and female (B) mice. Graphs on the left show mouse grouping for components 1 and 2 of the PLS regression. Individual mice are represented by symbols, and group SD by ellipsoids. Variance explained by each component is shown in parenthesis. Graphs on the right show fold-change of gene expression, and VIP scores calculated from the resulting PLS model. Filled symbols represent VIP>1.

Notably, HFHSD feeding for 3 days triggered different inflammatory responses in male and female mice. Namely, male mice showed increased expression of IL-13, IL18, IFN-γ, CD68 and EMR1 and a small decrease of IL-10 (figure 2A), while female mice had increased expression of IL-6 and IBA1, and reduced expression of IL-13 (figure 2B).

### Cytokine expression after 24 weeks of HFHSD-feeding and diet normalization

We next analyzed mice hypothalamic cytokine expression after 24 weeks of HFHSD. A PLS regression with 5 components provided a separation of mice in HFHSD or CD based on their gene expression (figure 3A-B) and explained 92% and 98% of the variance in males and females, respectively. Like 3 days of HFHSD, prolonged HFHSD feeding resulted in different cytokine expression profiles in each gender. Male mice presented an overall tendency for a decrease in pro- and anti-inflammatory cytokine mRNA expression after HFHSD, relative to CD, with important decreases in expression of TNF-α, IL-6 and IL-13, and an increase in TGF-β (figure 3A). In contrast, female mice on HFHSD showed generally unchanged cytokine expression, with increases in the expression of pro-inflammatory cytokines IL-18 and IFN-γ (figure 3B). The PLS regression models were then used to predict effects of diet normalization in mice of the RevD group. While the PLS component space indicates a cytokine expression shift in RevD mice, there was no full normalization of the cytokine profile in male (figure 3A) or female (figure 3B) mice. Only a few cytokines normalized their expression after diet reversal in male mice, such as TNF-α and IL-13 (figure 3C). Interestingly, while HFHSD-induced overexpression of IL-18 and IFN-γ were normalised in females, other cytokines appeared modified after diet reversal, namely TNFα (figure 3D).

**FIGURE 3.**
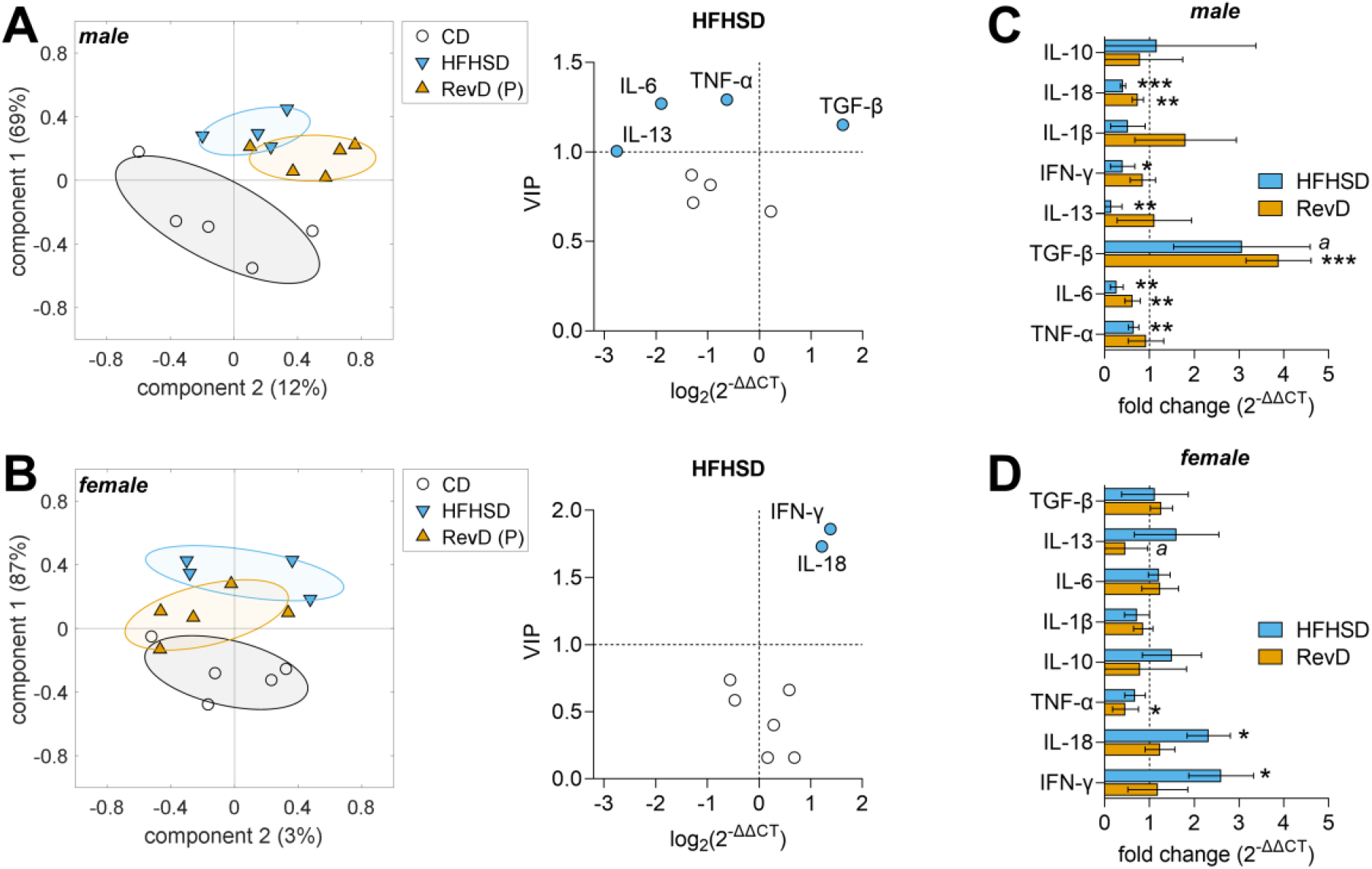
HFHSD feeding for 24 weeks induced gender-specific hypothalamic inflammation, that is, differential gene expression changes were observed in male (A) and female (B) mice. PLS regression models were constructed for HFHSD and CD, and then used to calculate the component space for predicting effects of diet normalization in RevD (P). Graphs on the left show mouse grouping for components 1 and 2 of the PLS regression. Individual mice are represented by symbols, and group SD by ellipsoids. Variance explained by each component is shown in parenthesis. Graphs on the right show fold-change of gene expression for HFHSD and RevD relative to CD, and VIP scores calculated from the resulting PLS model. Filled symbols represent VIP>1. Panels C and D show gene expression in HFHSD and RevD mice relative to control CD (mean±SD of n=5). Crescent VIP scores are represented from top to bottom. Significance for one sample t-test comparisons to 1 are indicated as follows *P<0.05, **P<0.01, ***P<0.001 (^a^ P=0.07).

### Gliosis in the hypothalamic arcuate nucleus

We next analyzed the morphology of astrocytes (GFAP^+^ cells) and microglia (IBA-1^+^ cells) in the arcuate nucleus (ARC) of hypothalamus by immunofluorescence imaging in mice fed with HFHSD for 24 weeks (figure 4A). We analyzed astrocytes by the number, total, and mean area of GFAP^+^ cells; and microglia by the total number and area of IBA1^+^ cells, as well as the percentage of activated microglia as morphologically defined elsewhere [30]. No major alterations were found between groups regarding GFAP immunostaining on cell number and area in male mice (figure 4B). In female mice, on the other hand, there was a tendency for increased GFAP area, which was normalized by diet reversal (figure 4C). While no major alterations were found between groups regarding total Iba1^+^ cell number and area, an increase on activated microglia with HFHSD was found in both male and female mice (P<0.05; figure 4B-C). While microglia remain activated when male mice were switched to control diet (figure 4B), microglia activation was reversed in females (P<0.01; figure 4C).

**FIGURE 4.**
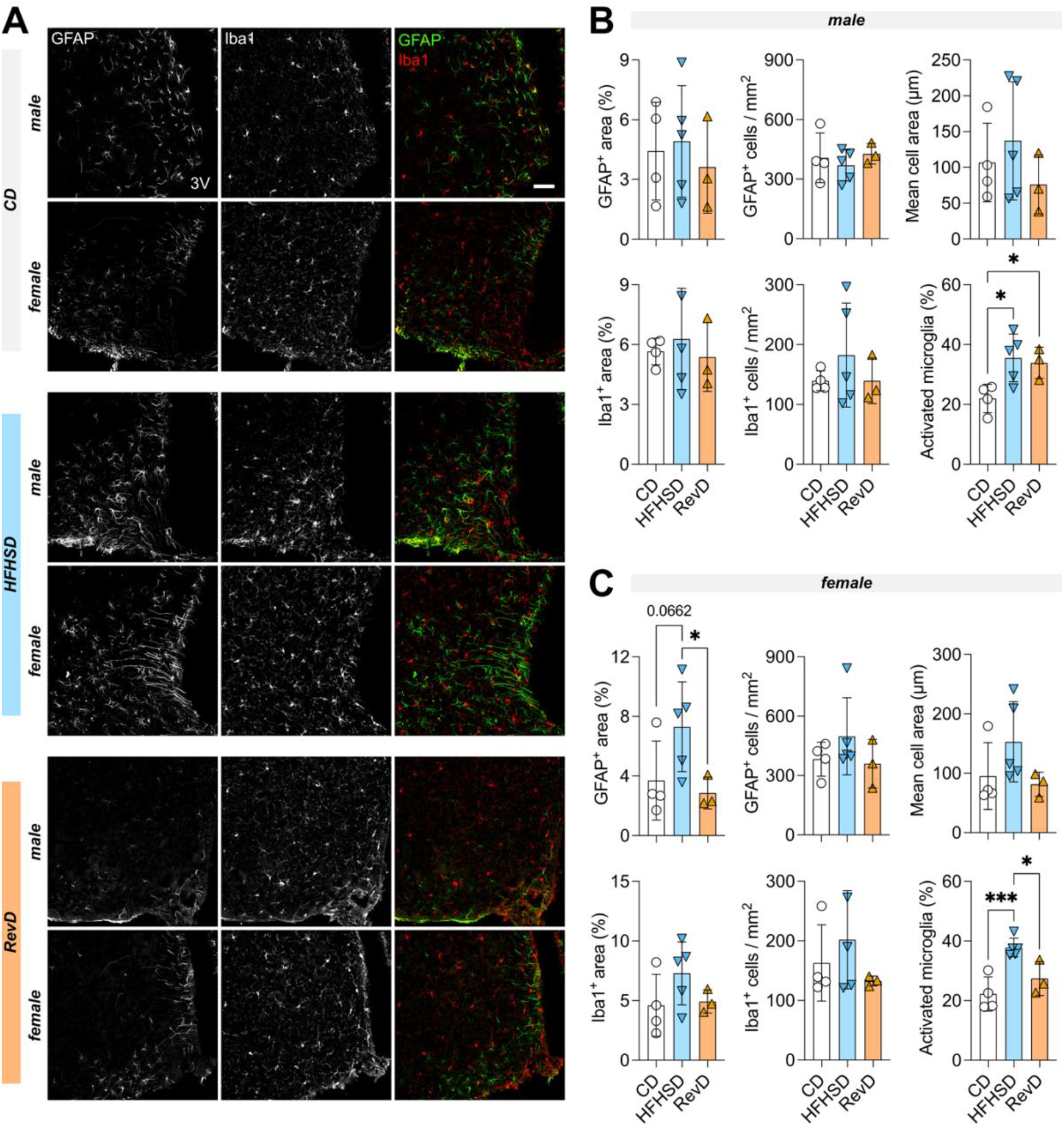
Astrogliosis and microgliosis in the arcuate nucleus of male and female mice. (A) representative micrographs of astrocytes (GFAP^+^ cells, green) and microglia (IBA1^+^ cells, red) cells in the arcuate nucleus (scale bar is 50 µm). Total GFAP stained area, number of GFAP^+^ cells or mean cell area were evaluated for astrogliosis while IBA1 stained area, number of IBA1^+^ cells or fraction of activated microglia (poorly ramified cells) were evaluated in male (B) and female (C) mice. Bars are mean±SD of n=4 (CD), n=5 (HFHSD), n=3 (RevD), and symbols represent individual mice. *p<0.05, ***P<0.001 from Fisher’s LSD test after significant ANOVA.

## DISCUSSION

Located in the mediobasal hypothalamus, the ARC is particularly positioned to sense circulating factors that regulate metabolism. Together with the median eminence, the ARC is a circumventricular organ that lacks a true blood–brain barrier (BBB) formed by endothelial cells, resulting in exposure to circulating factors [31]. Others have previously reported effects of obesogenic diets in the development of hypothalamic neuroinflammation [18-20, 22]. However, whether this is a sex-specific, reversible process still needs validation. In this study, we found that female mice fed HFHSD for 24 weeks exhibited a shift in cytokine mRNA levels, accompanied by astrocyte and microglia activation. These features were largely normalized to control levels after diet reversal. On the other hand, male mice presented an overall decrease in pro- and anti-inflammatory cytokine levels, which also tended to be normalized after diet reversal. Interestingly, female but not male mice showed astrogliosis after long-term HFHSD exposure. Since only females recovered microglia activation, we propose that astrogliosis in long-term HFHSD exposure is not triggering injury, but can rather facilitate the resolution of neuroinflammation. In contrast to our findings of astrogliosis in females but not males after obesogenic diet exposure for 24 weeks, a shorter 8-week HFD exposure was found to induce astroglyosis in the arcuate nucleus of males but not females [32].

It has been previously reviewed that both basal body inflammatory and immune responses have sex-specific differences based on genetic mediators (such as sex chromosomes and microRNAs and long non-coding RNAs), hormonal mediators (estradiol, progesterone, and androgens), and environmental mediators (*e.g*. nutrition and microbiome) [33]. It was formerly found that both innate and adaptive responses are generally higher in females compared to males. Females tend to have greater antibody levels and responses, higher immunoglobulin levels and higher B cell numbers [34, 35]. For example, it has been shown that female adult mice possess higher levels of T helper 1 cytokine producing cells, responsible for the production of INF-γ [36], which corroborates our findings. This gender-specific differences can be due to the fact that both androgen and estrogen response elements can be found in the promoters of several innate immunity genes, leading to a dimorphic immune response [37]. It was already shown that low levels of the female sex hormone estradiol can increase the production of pro-inflammatory cytokines IL-1, IL-6 and TNF-α, while higher levels of this hormone have the opposite effect [38]. On the other hand, male sex hormones androgens have been described to exhibited anti-inflammatory properties. Testosterone was revealed to increase the levels of anti-inflammatory cytokine TGF-β [39], while reducing the levels of pro-inflammatory cytokine TNF-α [40], as observed in the present study.

Few preclinical studies have looked at the effect of obesogenic diets comparing male and female, and even fewer took this gender consideration regarding hypothalamic inflammation. Daly *et al*. investigated sex differences when mice were fed with HFHSD and found that male mice displayed lower levels of the pro-inflammatory cytokines IL-1β and IL-6 compared sex-related mice fed a low-fat, low-sucrose diet (LFLSD) [26]. Contrarily, HFHSD fed female mice evidenced an increase in cytokines levels compared to the correspondent LFLSD group [40]. The authors found no alterations on TGF-β and TNF-α protein levels. Recently, a broad study comparing sex- and age-dependent behavior and inflammatory parameters in mouse under high-fat but not high sucrose diet, described alterations in plasma of young female mice, while no effects after 5-6 months of HFD were observed on young male mice [41]. Increased pro-inflammatory cytokines and chemokines such as IL17A/CTLA8, Eotaxin/CCL11, MCP3/CCL7, and Leptin were observed in HFD-fed females, with decreased levels of IL22 (the IL-10 family cytokine that is produced by T cells [41]. In the brain innate immunity, a decrease in microglial cell complexity in HFHSD male mice was found by Daly *et al*., a marker for cell activation. No changes were observed between the female mice groups. Interestingly, the same study reported major differences in gut microbiome species between all the different groups, and, more specifically, HFHSD male mice develop an increase gut microbiota species diversity compared to LFLSD. Moreover, a correlation between diet-induced gut microbiome alterations and hypothalamic inflammatory profile was evident [26]. This dysbiosis was previously shown to affect the central nervous system physiology and inflammation through the gut-brain axis, that encompasses a panoply of intricate pathways that include the vagal nerve, the immune system, and bacterial-derived metabolites [42, 43]. Since neuroinflammation can be a direct response to how components of the diet are metabolized after digestion and that many metabolites can specifically arise from gut microbiome metabolism, one can speculate that gut dysbiosis can be a major player in diet-induced neuroinflammation. Intestinal inflammation and increased permeability develop in adult male mice after 12 weeks of HFD or high sugar diet [44], with a clearly variable diet-dependent changes in the levels of cytokines in the colon of mice [44, 25]. However, more studies are needed on the sex-related alterations across the gut-brain axis and its connection between gut dysbiosis as a cause for neuroinflammation. Moreover, little is known about the effects of a RevD on the gut microbiome.

To our surprise, neuroinflammatory markers measured in our study were not strikingly increased after 3 days of HFHSD exposure, in contrast to observations by Thaler *et al*. using HFD [18]. Aside any possible experimental peculiarities on mouse strain, diet, housing or handling, our experience feeding obesogenic diets [14, 22, 23, 45] to mice leads us to believe that HFD alone might be a stronger inducer of metabolic syndrome than HFHSD. The lower severity of metabolic syndrome during HFHSD than during HFD is likely the reason for the present study to not reproduce the early hypothalamic inflammation reported for HFD-fed mice [18, 22, 23].

To conclude, mice fed HFHSD display complex sex-specific changes of inflammatory cytokine profiles in the hypothalamus that can be partially reversed by diet normalization. These cytokine changes are, however, not necessarily accompanied by or indicative of gliosis. In fact, male mice showed activation of microglia but not astrocytes upon HFHSD feeding, while female mice showed activation of both, and gliosis was reversible in females but not males.

## DATA AVAILABILITY STATEMENT

All data are contained within the manuscript and can be shared upon request to the corresponding author.

## ETHICS STATEMENT

### AUTHOR CONTRIBUTIONS

JMND designed the study. GCdP, RFS and AMGS and performed experiments and analyzed data. GCdP and RFS wrote the manuscript. All authors revised the manuscript.

## FUNDING

This work was supported by the Swedish foundation for International Cooperation in Research and Higher education (BR2019-8508), Swedish Research council (2019-01130), Diabetesfonden (DIA2019-440, DIA2021-637), and Direktör Albert Påhlssons Foundation. R.F.S. was funded by Tage Blücher Foundation. J.M.N.D. acknowledges generous financial support from The Knut and Alice Wallenberg foundation, the Faculty of Medicine at Lund University and Region Skåne. The authors acknowledge support from the Lund University Diabetes Centre, which is funded by the Swedish Research Council (Strategic Research Area EXODIAB, grant 2009-1039) and the Swedish Foundation for Strategic Research (grant IRC15-0067).

## ACKNOWLEDGMENTS

The Lund University Bioimaging Centre is acknowledged for access to microscopy resources.

## Notes

### Competing Interest Statement

The authors have declared no competing interest.

## REFERENCES

1. Collaboration, N.R.F., Trends in adult body-mass index in 200 countries from 1975 to 2014: a pooled analysis of 1698 population-based measurement studies with 19.2 million participants. Lancet, 2016. 387(10026): p. 1377–1396.

2. Wong, M.C.S., et al., Global, regional and time-trend prevalence of central obesity: a systematic review and meta-analysis of 13.2 million subjects. Eur J Epidemiol, 2020. 35(7): p. 673–683. 10.1007/s10654-020-00650-3

3. González-Muniesa, P., Mártinez-González, M. A., Hu, et al., Obesity. Nat Rev Dis Primers, 2017. 3: p. 17034. 10.1038/nrdp.2017.34

4. Williams, E.P., Mesidor, M., Winters, K., Dubbert, P. M., & Wyatt, S. B. Overweight and Obesity: Prevalence, Consequences, and Causes of a Growing Public Health Problem. Curr Obes Rep, 2015. 4(3): p. 363–70. 10.1007/s13679-015-0169-4

5. McAllister, E.J., Dhurandhar, N. V., Keith, S. W., et al., Ten putative contributors to the obesity epidemic. Crit Rev Food Sci Nutr, 2009. 49(10): p. 868–913. 10.1080/10408390903372599

6. Saeed, S., Bonnefond, A., Manzoor, J., Shabbir, F., et al., Genetic variants in LEP, LEPR, and MC4R explain 30% of severe obesity in children from a consanguineous population. Obesity (Silver Spring), 2015. 23(8): p. 1687–95. 10.1002/oby.21142

7. Ludwig, D.S., Lifespan Weighed Down by Diet. JAMA, 2016. 315(21): p. 2269–70. 10.1001/jama.2016.3829

8. Ng, M., Fleming, T., Robinson, M.,et al., Global, regional, and national prevalence of overweight and obesity in children and adults during 1980-2013: a systematic analysis for the Global Burden of Disease Study 2013. Lancet, 2014. 384(9945): p. 766–81. 10.1016/S0140-6736(14)60460-8

9. Bo, S., Gentile, L., Ciccone, G., et al., The metabolic syndrome and high C-reactive protein: prevalence and differences by sex in a southern-European population-based cohort. Diabetes Metab Res Rev, 2005. 21(6): p. 515–24. 10.1002/dmrr.561

10. Laaksonen, D.E., Niskanen, L., Punnonen, K., et al., Sex hormones, inflammation and the metabolic syndrome: a population-based study. Eur J Endocrinol, 2003. 149(6): p. 601–8. 10.1530/eje.0.1490601

11. Hotamisligil, G.S., Inflammation and metabolic disorders. Nature, 2006. 444(7121): p. 860–7. 10.1038/nature05485

12. Frisardi, V., Solfrizzi, V., Seripa, D., et al., Metabolic-cognitive syndrome: a cross-talk between metabolic syndrome and Alzheimer’s disease. Ageing Res Rev, 2010. 9(4): p. 399–417. 10.1016/j.arr.2010.04.007

13. Askari, M., Heshmati, J., Shahinfar, H., et al., Ultra-processed food and the risk of overweight and obesity: a systematic review and meta-analysis of observational studies. Int J Obes (Lond), 2020. 44(10): p. 2080–2091. 10.1038/s41366-020-00650-z

14. Garcia-Serrano, A.M., Mohr, A. A., Philippe, J., et al., Cognitive Impairment and Metabolite Profile Alterations in the Hippocampus and Cortex of Male and Female Mice Exposed to a Fat and Sugar-Rich Diet are Normalized by Diet Reversal. Aging Dis, 2022. 13(1): p. 267–283. 10.14336/AD.2021.0720

15. Williams, K.W. and J.K. Elmquist, From neuroanatomy to behavior: central integration of peripheral signals regulating feeding behavior. Nat Neurosci, 2012. 15(10): p. 1350–5. 10.1038/nn.3217

16. Konner, A.C., T. Klockener, and J.C. Bruning, Control of energy homeostasis by insulin and leptin: targeting the arcuate nucleus and beyond. Physiol Behav, 2009. 97(5): p. 632–8. 10.1016/j.physbeh.2009.03.027

17. Blouet, C. and G.J. Schwartz, Hypothalamic nutrient sensing in the control of energy homeostasis. Behav Brain Res, 2010. 209(1): p. 1–12. 10.1016/j.bbr.2009.12.024

18. Thaler, J.P., Yi, C. X., Schur, E. A., et al., Obesity is associated with hypothalamic injury in rodents and humans. J Clin Invest, 2012. 122(1): p. 153–62. 10.1172/JCI59660

19. Zhang, X., Zhang, G., Zhang, H. et al., Hypothalamic IKKbeta/NF-kappaB and ER stress link overnutrition to energy imbalance and obesity. Cell, 2008. 135(1): p. 61–73. 10.1016/j.cell.2008.07.043

20. De Souza, C.T., Araujo, E. P., Bordin, S., et al., Consumption of a fat-rich diet activates a proinflammatory response and induces insulin resistance in the hypothalamus. Endocrinology, 2005. 146(10): p. 4192–9. 10.1210/en.2004-1520

21. Mohr, A.A., Garcia-Serrano, A. M., Vieira, J. P., et al., A glucose-stimulated BOLD fMRI study of hypothalamic dysfunction in mice fed a high-fat and high-sucrose diet. J Cereb Blood Flow Metab, 2021. 41(7): p. 1734–1743. 10.1177/0271678X20942397

22. Lizarbe, B., Cherix, A., Duarte, J. M. N., et al., High-fat diet consumption alters energy metabolism in the mouse hypothalamus. Int J Obes (Lond), 2019. 43(6): p. 1295–1304.

23. Lizarbe, B., Soares, A. F., Larsson, S., Duarte, J. M. Neurochemical Modifications in the Hippocampus, Cortex and Hypothalamus of Mice Exposed to Long-Term High-Fat Diet. Front Neurosci, 2018. 12: p. 985. 10.3389/fnins.2018.00985

24. Casimiro, I., et al., Phenotypic sexual dimorphism in response to dietary fat manipulation in C57BL/6J mice. J Diabetes Complications, 2021. 35(2): p. 107795.

25. Church JS, Renzelman ML, Schwartzer JJ. Ten-week high fat and high sugar diets in mice alter gut-brain axis cytokines in a sex-dependent manner. J Nutr Biochem, 2022 Feb;100:108903. 10.1016/j.jnutbio.2021.108903

26. Daly, C.M., Saxena, J., Singh, J., et al., Sex differences in response to a high fat, high sucrose diet in both the gut microbiome and hypothalamic astrocytes and microglia. Nutr Neurosci, 2022. 25(2): p. 321–335. 10.1080/1028415X.2020.1752996

27. Dearden L, Balthasar N. Sexual dimorphism in offspring glucose-sensitive hypothalamic gene expression and physiological responses to maternal high-fat diet feeding. Endocrinology, 2014. 155(6):2144-54

28. Skoug, C., C. Holm, and J.M.N. Duarte, Hormone-sensitive lipase is localized at synapses and is necessary for normal memory functioning in mice. J Lipid Res, 2022. 63(5): p. 100195. 10.1016/j.jlr.2022.100195

29. Duarte, J.M., et al., Caffeine consumption prevents diabetes-induced memory impairment and synaptotoxicity in the hippocampus of NONcZNO10/LTJ mice. PLoS One, 2012. 7(4): p. e21899. 10.1371/journal.pone.0021899

30. Fernandez-Arjona, M.D.M., et al., Microglia Morphological Categorization in a Rat Model of Neuroinflammation by Hierarchical Cluster and Principal Components Analysis. Front Cell Neurosci, 2017. 11: p. 235. 10.3389/fncel.2017.00235

31. Rodríguez, E.M., Blázquez, J.L. & Guerra, M. The design of barriers in the hypothalamus allows the median eminence and the arcuate nucleus to enjoy private milieus: the former opens to the portal blood and the latter to the cerebrospinal fluid. Peptides, 2010. 31(4):757–76. 10.1016/j.peptides.2010.01.003

32. Morselli E, Fuente-Martin E, Finan B, et al., Hypothalamic PGC-1α protects against high-fat diet exposure by regulating ERα. Cell Rep, 2014. 9(2):633-45. 10.1016/j.celrep.2014.09.025

33. Klein, S.L. and K.L. Flanagan, Sex differences in immune responses. Nat Rev Immunol, 2016. 16(10): p. 626–38. 10.1038/nri.2016.90

34. Abdullah, M., Chai, P. S., Chong, M. Y., et al., Gender effect on in vitro lymphocyte subset levels of healthy individuals. Cell Immunol, 2012. 272(2): p. 214–9. 10.1016/j.cellimm.2011.10.009

35. Furman, D., et al., Systems analysis of sex differences reveals an immunosuppressive role for testosterone in the response to influenza vaccination. Proc Natl Acad Sci U S A, 2014. 111(2): p. 869–74. 10.1073/pnas.1321060111

36. Roberts, C.W., W. Walker, and J. Alexander, Sex-associated hormones and immunity to protozoan parasites. Clin Microbiol Rev, 2001. 14(3): p. 476–88. 10.1128/CMR.14.3.476-488.2001

37. Hannah, M.F., V.B. Bajic, and S.L. Klein, Sex differences in the recognition of and innate antiviral responses to Seoul virus in Norway rats. Brain Behav Immun, 2008. 22(4): p. 503–16. 10.1016/j.bbi.2007.10.005

38. Bouman, A., M.J. Heineman, and M.M. Faas, Sex hormones and the immune response in humans. Hum Reprod Update, 2005. 11(4): p. 411–23. 10.1093/humupd/dmi008

39. Liva, S.M. and R.R. Voskuhl, Testosterone acts directly on CD4+ T lymphocytes to increase IL-10 production. J Immunol, 2001. 167(4): p. 2060–7. 10.4049/jimmunol.167.4.2060

40. D’Agostino, P., Milano, S., Barbera, C., et al., Sex hormones modulate inflammatory mediators produced by macrophages. Ann N Y Acad Sci, 1999. 876: p. 426–9. 10.1111/j.1749-6632.1999.tb07667.x

41. Evans, A.K., Saw, N.L., Woods, C.E., et al, Impact of high-fat diet on cognitive behavior and central and systemic inflammation with aging and sex differences in mice. Brain, behavior, and immunity, 2024. 118, 334–354. 10.1016/j.bbi.2024.02.025

42. Burberry, A., Wells, M.F., Limone, F., et al., C9orf72 suppresses systemic and neural inflammation induced by gut bacteria. Nature, 2020. 582(7810): p. 89–94. 10.1038/s41586-020-2288-7

43. Rutsch, A., J.B. Kantsjo, and F. Ronchi, The Gut-Brain Axis: How Microbiota and Host Inflammasome Influence Brain Physiology and Pathology. Front Immunol, 2020. 11: p. 604179. 10.3389/fimmu.2020.604179

44. Do MH, Lee E, Oh M-J, Kim Y, Park H-Y. High-Glucose or -Fructose Diet Cause Changes of the Gut Microbiota and Metabolic Disorders in Mice without Body Weight Change. Nutrients, 2018; 10(6):761. 10.3390/nu10060761

45. Garcia-Serrano, A.M., Vieira, J.P.P., Fleischhart, V., Duarte, J.M.N., Taurine or N-acetylcysteine treatments prevent memory impairment and metabolite profile alterations in the hippocampus of high-fat diet-fed female mice. 2022: p. 2022.02.02.478774. 10.1080/1028415X.2022.2131062

